# Variation in predicted COVID-19 risk among lemurs and lorises

**DOI:** 10.1101/2021.02.03.429540

**Authors:** Amanda D. Melin, Joseph D. Orkin, Mareike C. Janiak, Alejandro Valenzuela, Lukas Kuderna, Frank Marrone, Hasinala Ramangason, Julie E. Horvath, Christian Roos, Andrew C. Kitchener, Chiea Chuen Khor, Weng Khong Lim, Jessica G. H. Lee, Patrick Tan, Govindhaswamy Umapathy, Muthuswamy Raveendran, R. Alan Harris, Ivo Gut, Marta Gut, Esther Lizano, Tilo Nadler, Dietmar Zinner, Steig E. Johnson, Erich D. Jarvis, Olivier Fedrigo, Dongdong Wu, Guojie Zhang, Kyle Kai-How Farh, Jeffrey Rogers, Tomas Marques-Bonet, Arcadi Navarro, David Juan, Paramjit S. Arora, James P. Higham

**Affiliations:** Department of Anthropology and Archaeology, University of Calgary, Canada; Department of Medical Genetics, University of Calgary, Canada; Alberta Children’s Hospital Research Institute, University of Calgary, Canada; Experimental and Health Sciences Department (DCEXS), Institut de Biologia Evolutiva, Universitat Pompeu Fabra-CSIC, Barcelona, Spain; School of Science, Engineering & Environment, University of Salford, United Kingdom; Department of Chemistry, New York University, United States; Genomics & Microbiology Research Laboratory, North Carolina Museum of Natural Sciences, Raleigh, NC, USA; Department of Biological and Biomedical Sciences, North Carolina Central University, Durham, NC, USA; Department of Evolutionary Anthropology, Duke University, Durham, NC, USA; Department of Biological Sciences, North Carolina State University, Raleigh, NC, USA; Gene Bank of Primates and Primate Genetics Laboratory, German Primate Center, Leibniz Institute for Primate Research, Göettingen, Germany; Department of Natural Sciences, National Museums Scotland and School of Geosciences, University of Edinburgh, Edinburgh, United Kingdom; Genome Institute of Singapore, Agency for Science, Technology and Research, Singapore; Singapore Eye Research Institute, Singapore National Eye Centre, Singapore; SingHealth Duke-NUS Institute of Precision Medicine, Singapore Health Services, Singapore; SingHealth Duke-NUS Genomic Medicine Centre, Singapore Health Services, Singapore; Cancer and Stem Cell Biology Program, Duke-NUS Medical School, Singapore; Department of Conservation, Research and Veterinary Services, Wildlife Reserves Singapore, Singapore; CSIR-Laboratory for the Conservation of Endangered Species, Centre for Cellular and Molecular Biology, Hyderabad, India; Human Genome Sequencing Center and Department of Molecular and Human Genetics, Baylor College of Medicine, Houston, Texas, United States; Universitat Pompeu Fabra (UPF), Barcelona, Spain; Cuc Phuong Commune, Nho Quan District, Ninh Binh Province, Vietnam; Cognitive Ethology Laboratory, German Primate Center, Leibniz Institute for Primate Research, Goettingen, Germany; Leibniz Science Campus Primate Cognition, Goettingen, Germany; Department of Primate Cognition, Georg-August-University, Goettingen, Germany; The Vertebrate Genomes Lab, The Rockefeller University, New York, United States; Laboratory of Neurogenetics of Language, The Rockefeller University, New York, United States; Howard Hughes Medical Institute, Chevy Chase, Maryland, United States; State Key Laboratory of Genetic Resources and Evolution, Kunming Institute of Zoology, Chinese Academy of Sciences, Kunming, China; Kunming Natural History Museum of Zoology, Kunming Institute of Zoology, Chinese Academy of Sciences, Kunming, China; Villum Center for Biodiversity Genomics, Section for Ecology and Evolution, Department of Biology, University of Copenhagen, Denmark; China National Genebank, BGI-Shenzhen, Shenzhen, 518083, China; Center for Excellence in Animal Evolution and Genetics, Chinese Academy of Sciences, Kunming, China; Artificial Intelligence Lab, Illumina Inc, San Diego, CA, USA; Catalan Institution of Research and Advanced Studies (ICREA), Barcelona, Spain; CNAG-CRG, Centre for Genomic Regulation (CRG), Barcelona Institute of Science and Technology (BIST), Barcelona, Spain; Institut Català de Paleontologia Miquel Crusafont, Universitat Autònoma de Barcelona, Barcelona, Spain; Department of Anthropology, New York University, United States; New York Consortium in Evolutionary Primatology, New York, United States

## Abstract

The novel coronavirus SARS-CoV-2, which in humans leads to the disease COVID-19, has caused global disruption and more than 1.5 million fatalities since it first emerged in late 2019. As we write, infection rates are currently at their highest point globally and are rising extremely rapidly in some areas due to more infectious variants. The primary viral target is the cellular receptor angiotensin-converting enzyme-2 (ACE2). Recent sequence analyses of the *ACE2* gene predicts that many nonhuman primates are also likely to be highly susceptible to infection. However, the anticipated risk is not equal across the Order. Furthermore, some taxonomic groups show high ACE2 amino acid conservation, while others exhibit high variability at this locus. As an example of the latter, analyses of strepsirrhine primate *ACE2* sequences to date indicate large variation among lemurs and lorises compared to other primate clades despite low sampling effort. Here, we report *ACE2* gene and protein sequences for 71 individual strepsirrhines, spanning 51 species and 19 genera. Our study reinforces previous results and finds additional variability in other strepsirrhine species, and suggests several clades of lemurs have high potential susceptibility to SARS-CoV-2 infection. Troublingly, some species, including the rare and Endangered aye-aye (*Daubentonia madagascariensis*), as well as those in the genera *Avahi* and *Propithecus*, may be at high risk. Given that lemurs are endemic to Madagascar and among the primates at highest risk of extinction globally, further understanding of the potential threat of COVID-19 to their health should be a conservation priority. All feasible actions should be taken to limit their exposure to SARS-CoV-2.

## Introduction

On Tuesday September 22, 2020, the one-millionth human death officially attributed to COVID-19 was documented by the Johns Hopkins University Coronavirus Resource Center (BBC News, 2020). Since this date the rates of infection by the virus responsible for this disease, SARS-CoV-2, have increased in most countries. As we write, we are reaching new global highs in active cases and witnessing the spread of new, more transmissible variants (Mahase, 2020; World Health Organization, 2020). As coordinated efforts within and across institutions, countries, and continents seek to identify treatments, develop vaccines, and curb the spread of this highly contagious virus, attention has also turned to the potential risks posed to nonhuman species (Damas et al., 2020; Liu et al., 2020; Melin et al., 2020; Wu et al., 2020). Zoonotic transfer of diseases from humans to nonhuman primates poses a major risk given the many physiological and genetic similarities shared within the Order Primates, and is a potentially grave risk to already endangered and fragmented populations (Gillespie & Leendertz, 2020).

In recent studies, the susceptibility of primates and other mammals to potential SARS-CoV-2 infection has been assessed by analysis of the gene sequences that code for the primary viral target, angiotensin-converting enzyme-2 (ACE2; (Damas et al., 2020; Delgado Blanco et al., 2020; Liu et al., 2020; Melin et al., 2020)). Receptor-virus interaction models, including by members of the present authorship, have highlighted the likely susceptibility of many species, especially of apes and monkeys of Africa and Asia (Parvorder Catarrhini); meanwhile, monkeys in the Americas (Parvorder Platyrrhini) are predicted to exhibit lower susceptibility (Liu et al., 2020; Melin et al., 2020). One striking feature of these analyses is the uniformity within these parvorders. Across the identified primary viral binding sites, all catarrhines exhibit one set of amino acid residues, and all platyrrhines another. Surprisingly, although gene sequences are only publicly available for a few strepsirrhine species (four lemurs and one galago), ACE2 variation in that suborder far exceeds variation present in the rest of the primate taxa examined to date (24 species spanning 21 genera and including tarsiers (1), platyrrhines (6), and catarrhines (14)). Of particular concern is the high sequence similarity at binding sites to human ACE2 of some lemur ACE2 proteins, including aye-ayes (*Daubentonia madagascariensis*) and Coquerel’s sifakas (*Propithecus coquereli*), which exhibited residues that are far more similar to those of humans and other catarrhines than to those present in platyrrhines (monkeys of the Americas) (Melin et al., 2020).

These findings raise questions about the variability and molecular evolution of the *ACE2* gene across Strepsirrhini, as well as about the potential susceptibility to initial infection by SARS-CoV-2 of different species across the suborder. Here, we expand substantially on previous reports of strepsirrhine ACE2 variation (Damas et al., 2020; Melin et al., 2020). We report 71 *ACE2* gene sequences, including 66 from unpublished strepsirrhine genomes, spanning 39 lemuriform and 12 lorisiform species. For residue variants that have not been previously identified and assessed, we model the interactions between the translated ACE2 protein and the receptor-binding domain of the SARS-CoV-2 spike protein to predict the susceptibility of species to initial infection by SARS-CoV-2. In doing so, we seek to improve our understanding of ACE2 variation and evolution, and to identify which strepsirrhine species are likely to be most at risk from the COVID-19 pandemic. We recognize that disease development, progression and pathogenesis in any given species will also be impacted by factors influencing the efficacy of viral cellular entry and taxon-specific immune responses (Hoffmann et al., 2020; Li et al., 2020; Lukassen et al., 2020). Nonetheless, our hope is that our analysis of the initial susceptibility of different lemur and loris species to SARS-CoV-2 infection will help to inform decisions on how best to proceed with strepsirrhine research and management programs.

## Methods

### Study Species

We examined the *ACE2* gene sequence of 71 individual strepsirrhines - 66 newly sequenced individuals as part of the Primate Variation Genome Consortium (in preparation, Supple. Table 1), plus five obtained from publicly available genomes ^6^: *Otolemur garnettii* (Northern greater galago), accession no: XM_003791864.2, gene ID: 100951881; *Propithecus coquereli* (Coquerel’s sifaka), accession no: XM_012638731.1, gene ID: 105805773; *Microcebus murinus* (gray mouse lemur) accession no: XM_020285237.1, gene ID: 105882317; *Eulemur flavifrons* (blue-eyed black lemur), accession no: LGHW01000591.1, scaffold 590 (gene identified via BLAST); *Daubentonia madagascariensis*, (aye-aye) accession no: PVJZ01006595.1, scaffold 13170, (gene identified via BLAST). In total, we analyze the *ACE2* gene and protein sequences of 51 species (39 Lemuriformes and 12 Lorisiformes) and 19 genera (12 Lemuriformes and 7 Lorisiformes) (Table 1; Supple. Table 1). In addition, we model the impact of residues at binding sites recently reported for *Indri indri* ACE2 protein (Damas et al., 2020), adding another genus of lemur to our survey.

**Table 1.** Results of computational protein-protein interaction experiments predicting impact of amino acid changes, relative to human ACE2 residues, at critical binding sites with SARS-CoV-2 receptor binding domain. Impacts of changes across the full complement of critical binding sites are presented in (A), single residue replacements are presented in (B).

| Species | Mutations (relative to human sequence) | $\Delta\Delta G$ (kcal/mol)<sup>†</sup> | Estimated decrease in binding efficiency (log)<sup>‡</sup> |
| --- | --- | --- | --- |
| *Homo sapiens* |  | 0 |  |
| *Avahi, Daubentonia, Propithecus* | M82T | +0.9 | 5 fold |
| *Eulemur, Hapalemur, Lemur, Prolemur, Varecia* | Q24E, M82T | +1.4 | 10 fold |
| *Indri* | H34N, M82N | +2.7 | 100 fold |
| Monkeys (Americas) | Y41H, Q42E, M42T | +3.5 | 400 fold |
| *Galago, Loris, Nycticebus, Otolemur* | H34R, D38E, Y41H, M82T | +3.8 | 600 fold |
| *Lepilemur, Microcebus, Mirza* | D30E, H34N, Y41H, M82T | +4.0 | 800 fold |
| *Arctocebus, Perodicticus* | Q24H, H34R, D38E, Y41H, M82T | +4.2 | 1200 fold |
| *Galagoides* | H34R, D38E, Y41H, M82T, Y83F | +4.4 | 1700 fold |
| *Cheirogaleus* | D30E, H34N, Y41H, M82T, K353Q | +5.0 | 4500 fold |
| *Carlito* | H34Q, Y41H, M82S, K353N | +5.5 | 10000 |

**B)**
| Mutation | $\Delta\Delta G$ (kcal/mol)<sup>†</sup> |
| --- | --- |
| Q24L | -0.7 |
| Q24H | 0.3 |
| Q24E | 0.5 |
| D30E | 0.7 |
| H34R | 0.8 |
| H34N | 1.0 |
| H34L | -0.5 |
| H34Y | -0.1 |
| D38E | 0.6 |
| Y41H | 1.9 |
| Q42E | 0.9 |
| M82T | 0.9 |
| M82N | 1.8 |
| Y83F | 0.6 |
| K353Q | 0.9 |

<sup>†</sup>Approximate predicted decrease in binding affinity of individual ACE2 proteins and SARS-CoV-2 RBD. <sup>†</sup>Mutations were analyzed with SSIPe server (https://zhanglab.ccmb.med.umich.edu/SSIPe/) and PDB file 6M0J.

### Gene Alignments

We mapped reads from whole-genome sequence (WGS) data to the closest available annotated reference assembly from among the set of partially unpublished (unp.) references (*Daubentonia madagascariensis*, (unp.) *Galago moholi* (unp.), *Propithecus coquereli* (GCF_000956105.1), *Lemur catta* (unp.), *Loris tardigradus* (unp.), *Microcebus murinus* (unp.), *Nycticebus pygmaeus* (unp.) and *Otolemur garnettii* (unp.). Briefly, after removing adapter sequences using cutadapt, we mapped the reads using bwa mem and processed and sorted alignments using samtools. We removed duplicates using biobambam, and added read groups for variant calling using picard. We called variants using GATK4 HaplotypeCaller (v 4.1.6) following best practice pipelines (https://gatk.broadinstitute.org). After applying a set of standard hard filters, we extracted the coding regions of *ACE2* gene sequences and introduced homozygous alternative calls to create the putative coding sequence of each individual. We extracted and aligned *ACE2* gene sequences from the variant call files, which were then translated into protein sequences. The consensus sequences were manually inspected and corrected where needed to remove gene-flanking regions. We manually verified the absence of indels and premature stop codons for each individual. We then aligned these 66 amino acid sequences using MAFFT with default settings with those extracted from publicly available genomes (Melin et al., 2020). The full nucleotide and amino acid alignments used here are provided as a data table in the Supporting Information. Following alignment, we examined amino acid sequence variation within and across species along the length of the ACE2 protein, and specifically at the sites that are critical for SARS-CoV-2 binding.

### Variation in ACE2 sequences at critical sites and impact on SARS-CoV-2 binding

Our method for identifying critical contact sites between the ACE2 protein and the receptor-binding domain (RBD) of the SARS-CoV-2 spike protein is detailed in (Melin et al., 2020). Briefly, we conducted alanine scanning mutagenesis to assess the contribution of each human ACE2 residue to protein-protein complex formation with the SARS-CoV-2 RBD (Bogan & Thorn, 1998; Kortemme & Baker, 2002; Massova & Kollman, 1999). Alanine scanning is a commonly used method, and alanine is chosen because it is the smallest residue that may be incorporated without significantly impacting the protein backbone conformation (Kortemme et al., 2004). We defined critical residues as those that upon mutation to alanine decrease the binding energy by a threshold value ΔΔG_bind_ ≥1.0 kcal/mol. Nine sites meet this criterion (Supple. Table 2). To be conservative, we also examined amino acid variation at additional sites that were identified as important by different but complementary methods - cryo EM and X-ray crystallography structural analysis (Lan et al., 2020; Shang et al., 2020; Wang et al., 2020; Yan et al., 2020). While some of these sites overlap with the critical sites we identified using alanine scanning, three do not - alanine scanning also identifies these as binding sites, but with ΔΔG_bind_ <1.0 kcal/mol. For the present analyses, and as in Melin et al. 2020, we added these three sites suggested by other methods to be especially important binding sites to our nine sites for a total of 12 critical sites (Supple. Table 2). All computational alanine scanning mutagenesis analyses were performed using Rosetta (Jochim & Arora, 2010; Kortemme et al., 2004; Raj et al., 2013).

To model how variation in the ACE2 amino acid sequences across species affects the relative binding energy of the ACE2/SARS-CoV-2 interaction, we used the SSIPe program (Huang et al., 2019). This algorithm mutates selected residues and compares the resulting binding energy to that of human ACE2 bound to the SARS-CoV-2 RBD as a benchmark (PDB 6M0J). We modeled the full suite of amino acid changes occurring at critical binding sites for all unique ACE2 sequences. We further examined the predicted effect of each individual amino acid change (in isolation) on protein-binding affinity to better understand each residue’s contribution to variation in binding affinity.

## Results

### Variation in ACE2 sequences at critical sites and impact on SARS-CoV-2 binding

We examined variation along the length of the ACE2 protein sequence for the 66 newly sequenced individuals and the ACE2 sequences from the five genomes available at NCBI. Sequences are conserved within, but are variable across, strepsirrhine genera. The mean pairwise amino acid sequence identity along the length of the ACE2 protein within genera is 99.25% (mean amino acid substitutions = 5.85). The mean amino acid sequence identity between lemur genera was 91.67%, and between lorisiform genera was 92.72%. When we focused solely on the critical binding sites, the pairwise amino acid identity at binding sites within genera was 100%, indicating an absence of any amino acid variation among species in the same genus in our study. Differences between genera are also present at the critical binding sites (Figure 1), especially among lemurs, where the mean pairwise amino acid sequence identity is 83.18%. The mean pairwise amino acid sequence identity at binding sites between lorisiform genera is 95.04%.

**Figure 1.**
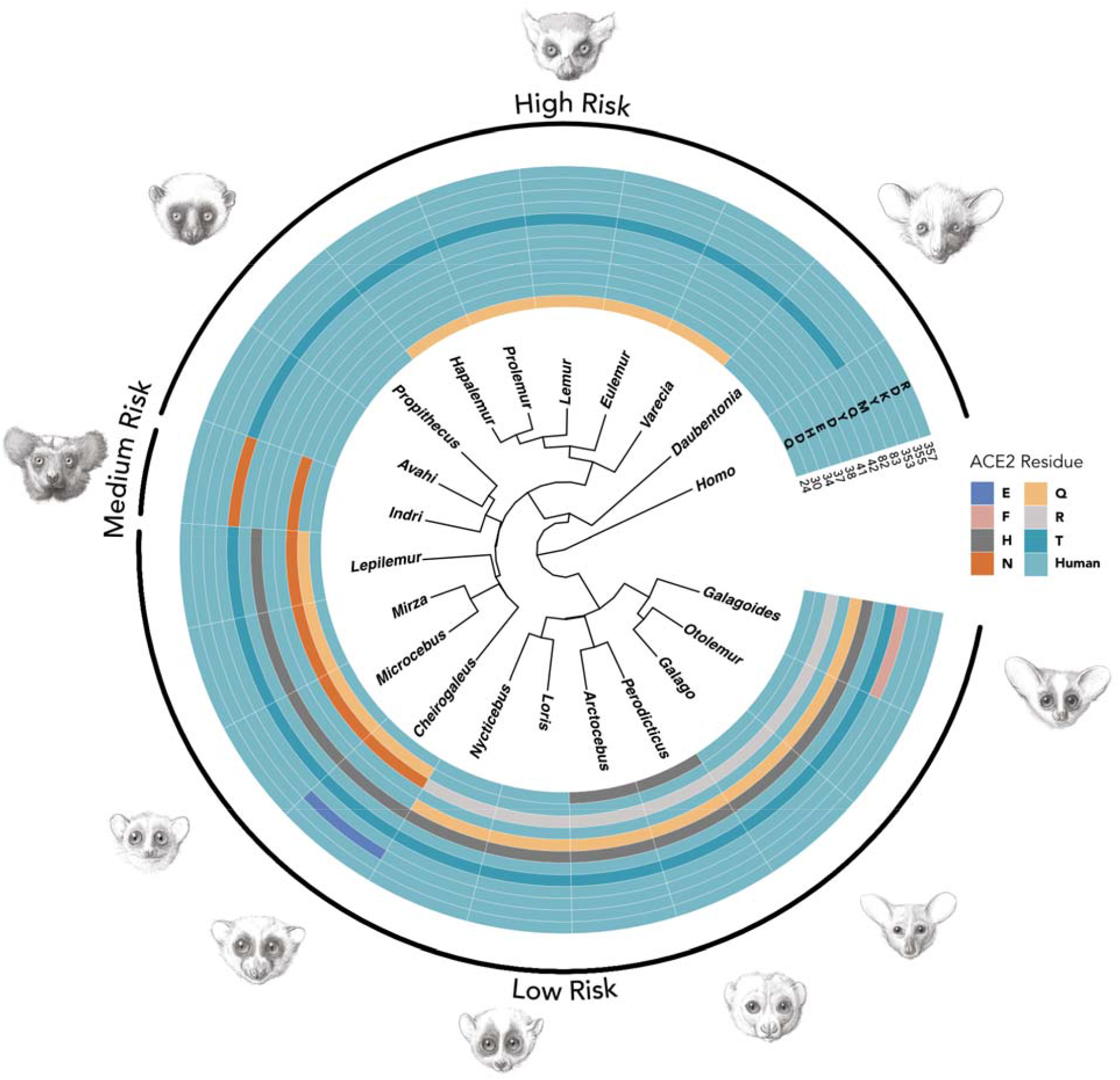
Amino acid composition of the ACE2 protein at sites that are critical for interacting with the SARS-CoV-2 receptor binding domain across examined strepsirrhine genera, with respect to the human ACE2 sequence.

**Figure 2.**
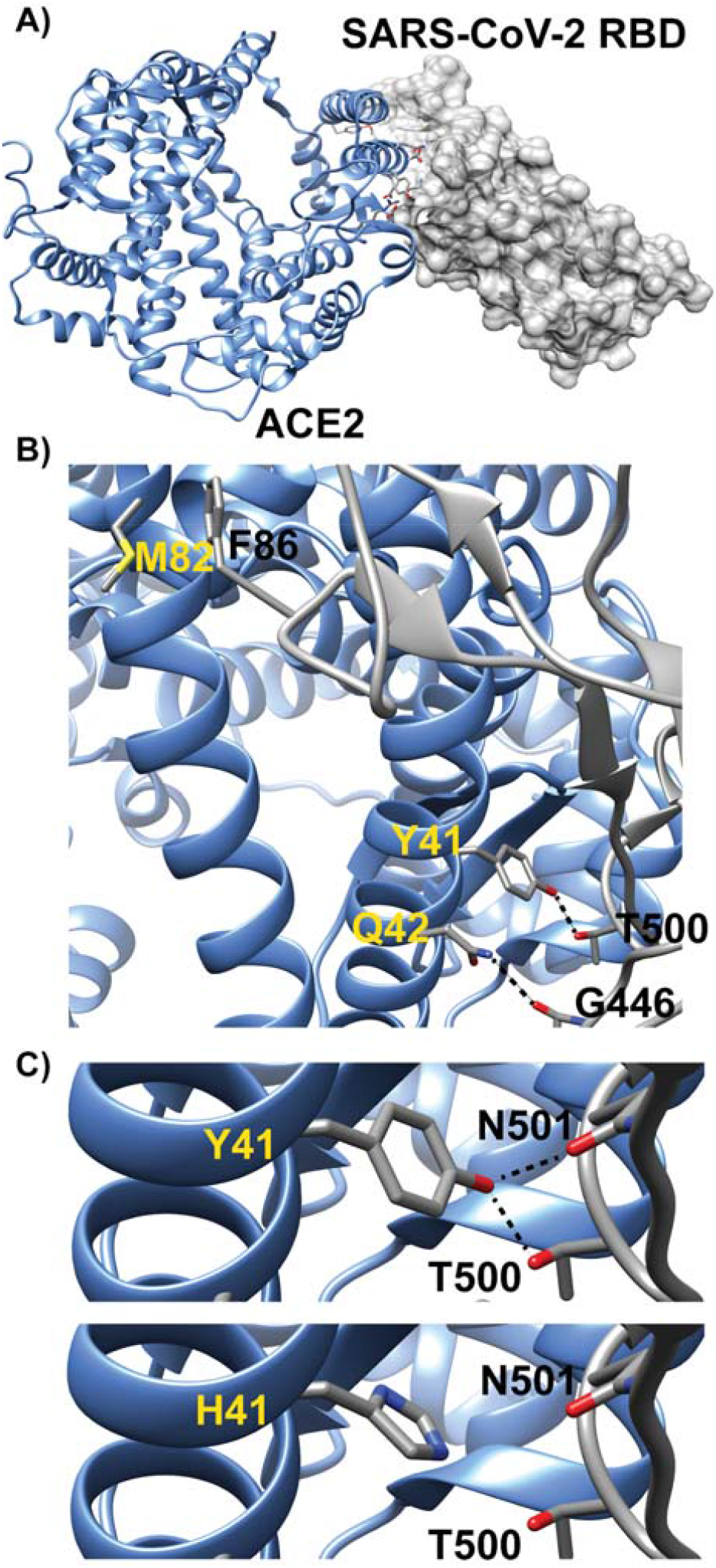
Model of human ACE2 in complex with SARS-CoV-2 RBD is presented in panel A. Three of the key ACE2 interfacial residues are highlighted in panel B. The dashed lines indicate predicted hydrogen bonding interactions. Interactions at three of the critical binding sites (41, 42, and 82) are shown for humans and other catarrhines (i.e. apes and monkeys in Africa and Asia). Changes in amino acids diminish these binding interactions. For example, substitution of tyrosine with histidine at site 41 in panel C decreases the viral-RBD binding affinity by removal of the potential hydrogen bonding interactions with T500 and N501.

We found three novel variants not previously reported for any primate at three critical binding sites: H24 (*Arctocebus, Perodicticus*), F83 (*Galagoides*), and Q353 (*Cheirogaleus*)). The remaining binding site variation is consistent with previous reports. The combination of residues (E24, T82) previously reported for *Eulemur flavifrons*, is also found in other species of *Eulemur*, as well as in *Hapalemur, Lemur*, and *Prolemur*. The critical binding site composition found in *Indri indri* (Damas et al., 2020) is not found in any of the genera we sequenced here. The residues at binding sites 37, 42, 355 and 357 are invariant across all strepsirrhines examined. None of the strepsirrhine ACE2 proteins are modeled to have higher binding affinity to SARS-CoV-2 than the human (catarrhine) form (Table 1A). Among the 12 critical sites, the substitutions causing the starkest drop in viral-receptor binding affinity relative to the human sequence are Y41H and M82N (Table 1B). The former substitution is found in all lorisoids and in sportive, dwarf, mouse, and giant mouse lemurs. The latter substitution is only identified in *Indri indri*, although a different substitution at the same site occurs in all other strepsirrhines (M82T); other mutations had lesser effects (Table 1B).

Looking at taxon-specific predictions based on the entire complement of amino acids at critical binding sites, the species predicted to be most susceptible to SARS-CoV-2 infection are in the genera *Avahi, Propithecus*, and *Daubentonia* (Table 1A). These taxa differ from humans at only one critical binding site, M82T, which is predicted to lower the binding affinity between the ACE2 receptor and the SARS-CoV-2 virus by 5-fold. These genera are followed by species in the genera *Eulemur, Lemur, Prolemur* and *Varecia*, which differ in one additional (Q24E) substitution, which should further lower the binding affinity by 2-fold. In potentially promising results, we predict that the lorises, galagos, and the dwarf, mouse, giant mouse, and sportive lemurs are far less susceptible to infection than humans. This is primarily due to a Y41H mutation, although additional changes in amino acids at binding sites further lower the affinity between their ACE2 and the RBD of the SARS-CoV-2 spike protein. The decreases in the modeled binding affinity range from 0.9 ΔΔG (kcal/mol) (*Avahi, Propithecus, Daubentonia;* predicted most susceptible) to 5.0 ΔΔG (kcal/mol)^a^ (*Cheirogaleus*, predicted least susceptible), relative to human ACE2 (Table 1A).

## Discussion

We report *ACE2* gene and protein sequences for 19 genera of strepsirrhine primates, spanning 51 species and 71 individuals, and examine these together with the *Indri indri* ACE2 protein. We confirm previous reports of the amino acid residue composition at viral binding sites for *Daubentonia, Propithecus, Eulemur, Microcebus*, and *Otolemur* (Damas et al., 2020; Melin et al., 2020). Additionally, we identified three novel variants at the following key binding sites: H24, F8, Q353. These variants are modeled to be protective and not found in species reported in the previous analyses of a small subset of strepsirrhine species (Melin et al. 2020), and were also not found among previous analyses of primates more generally. Relative to variation seen in catarrhines and platyrrhines, strepsirrhine ACE2 variation across genera at critical binding sites is remarkably high, especially among lemurs. In addition to reporting new ACE2 sequences spanning many strepsirrhine species, we also provide the first examination of intraspecific variation in ACE2 sequences outside of humans and vervet monkeys (Cao et al., 2020; Schmitt et al., 2020; Stawiski et al., 2020). We find that ACE2 proteins are highly conserved within species and within genera, at least for the taxa examined. Although our intraspecific and intrageneric sample sizes are small, this indicates that members of the same species and closely related species are likely to share similar initial susceptibility to SARS-CoV-2 infection. This allows predictions to be made regarding the likely susceptibility of species that are not included in our analyses, but that are closely-related to studied species.

As with all studies based on predictive modeling, our results should be interpreted with caution, especially those results which predict that some strepsirrhines might be at lower risk. Our results require experimental validation. Additional limitations include that our study examined variation at sites identified to be critical for SARS-CoV-2 viral binding, but did not assess the impact of residues that are not in direct contact with the virus and which may still affect binding allosterically. In addition, we did not examine genetic variation or model the function of the protease (TMPRSS2) that facilitates viral entry post binding (Hoffman et al. 2020), which is anticipated to impact disease progression. We also emphasize that our approach investigates the likely initial susceptibility of species to SARS-CoV-2 infection. The severity of viral infection responses may differ between species and is related to variation in immune and other responses (Lukassen et al., 2020). Nonetheless, the results of *in vivo* infection studies conducted on haplorrhine primates and other mammals strongly support the predictions of protein-protein interaction models about the susceptibility of different species to SARS-CoV-2 and the development of COVID-19-like symptoms (Blair et al., 2020; Lu et al., 2020; Rockx et al., 2020; Shan et al., 2020; Shi et al., 2020), supporting the applicability of our results. An additional tangible contribution of our study is that it provides novel sequence data that can be used in site-directed mutagenesis to recreate taxon-specific ACE2 proteins for cellular assays (Guy et al., 2005). At the same time, results predicting high susceptibility among a large number of genera are sufficiently alarming as to warrant special care and attention when interacting with these species in wild and captive management settings, including zoological parks, where humans frequently come into contact with vulnerable species (for example, in walk-through lemur exhibits).

The predictions of our study suggest that, among the strepsirrhines, the lemurs of the families Indriidae, Daubentoniidae, and to a lesser extent, Lemuridae are likely to be particularly vulnerable to SARS-CoV-2 infection. Lemurs are considered to be among the most threatened vertebrates globally, with over 94% of extant species being threatened with extinction (Schwitzer et al., 2014; IUCN, 2021). High rates of deforestation that result from changing land-use patterns coupled with high human population growth are among the most potent threats to lemur populations (Elmqvist et al., 2007; Harper et al., 2007). Besides habitat loss, these human-induced disturbances are exposing wild lemur populations to novel interactions with humans and domestic animals, and in so doing increasing risks of disease outbreaks (Barrett et al., 2013). Current knowledge on the disease ecology of wild lemurs suggests that populations that are found in disturbed habitats and/or those with a high volume of tourists are at elevated risk of bearing pathogens that are found in humans, livestock, and domestic animals (Bublitz et al., 2015; Junge, Barrett, and Yoder, 2011; Rasambainarivo et al., 2013). Furthermore, lemurs in general, including many of the larger bodied species that are predicted to be most at risk of infection by SARS-CoV-2, are highly vulnerable to new diseases because they are considered to be immunologically naïve, and unlikely to persist through a major epidemic outbreak (Junge 2007).

Historically, conservationists in Madagascar have strived to implement integrative conservation programs to protect the biological uniqueness of the island while leveraging the sustainable development of local communities (Corson, 2017). Conservation programs on the island implement strategies that include ecological research, monitoring, and restoration, agricultural intensification, alternative income sources such as ecotourism, and education (e.g., Wright et al., 2012; Birkinshaw et al., 2013; Dolins et al., 2010). In addition to the considerable conservation obstacles exacerbated by frequent political unrest and extreme poverty (Schwitzer et al., 2014), initiatives in Madagascar now have the concern of a viral pandemic in the human population, which appears likely to pose a direct and serious risk to many lemurs. Recently, there has been an emergence of integrated conservation programs that include a human health component e.g. (Mohan and Shellard, 2014; Garchitorena et al., 2018), a likely necessary shift in a country with one of the lowest levels of financing for the healthcare system (Barmania, 2015). This is particularly urgent given that COVID-19 could potentially infect up to 30% of the human population (Evans et al., 2020). Furthermore, Madagascar lacks a legal framework to guide best research practices to limit exposure of lemur population to diseases, in contrast, for example, to their great ape counterparts (see Gilardi et al. 2015). It is important that all stakeholders involved in the conservation of lemurs coordinate to draft such guidelines, which should include safeguards against relatively close contacts between people and lemurs that emerge as part of the successful and widespread community-based conservation programs. At a minimum these concerns should be carefully considered for the species identified as most susceptible to SARS-CoV-2.

In potentially more promising news, our results indicate that the vulnerability of the lorisoids – and indeed of many of the small-bodied lemurs, e.g. mouse lemurs – to SARS-CoV-2 infection may be substantially lower. Although the lorisoids as a group are more geographically widespread and (in relative terms) of lower conservation concern than the lemurs, many remain threatened and face critical challenges to their survival. Their cryptic, nocturnal habits may make the impact of emerging infectious diseases especially difficult to monitor in wild populations. Nonetheless, a lower predicted susceptibility to SARS-CoV-2 is a potentially positive result.

Our results are also likely to be of interest and significance for zoos and captive research facilities around the world that house lemurs, lorises, and galagos. Given the close contact with humans in such environments (often indoors), extra precautions may be necessary, especially when interacting with the likely most at-risk lemurs. These may include many of the measures suggested for those interacting with great ape and other catarrhine populations (Gilardi et al., 2015; Melin et al., 2020), such as regular testing for SARS-CoV-2, requiring face masks for human researchers and caretakers, imposing quarantines on all individuals ahead of contact, and disinfecting clothes and footwear. The risk of COVID-19 to nonhuman primates has been recently exemplified when members of a captive gorilla group tested positive for SARS-CoV-2 after exposure to an asymptomatic keeper (San Diego Zoo, 2021). Regardless of where species fall in the continuum of potential risk that we have presented here, we stress that it is prudent to take all feasible precautions when interacting with any primate.

Over the past year, scientists have mobilized with remarkable speed and effectiveness to address the COVID-19 pandemic’s impacts on humans, from developing and rolling out new tests, to designing and trialing vaccine candidates. The pandemic also raises major new challenges for field primatologists and other biologists, zookeepers, conservationists, and all those interested in the survival and welfare of primates (Douglas et al., 2020; Olival et al., 2020; Bales, 2020). Conservation efforts have already been severely impeded by the pandemic. National and global lock-downs have made it difficult or impossible for conservationists, primatologists, and wildlife patrol teams to enter their field sites. Governments are preoccupied with efforts to curb the pandemic, and in some cases conservation funding support has been reduced due to financial difficulties and new priorities. While avoiding all contact with wild primates may be ideal from a zoonotic disease containment perspective, it is unlikely to be possible in practice, nor in the overall interest of their conservation. While we hope that our results will serve to inform conservation efforts, resolving the pressures that have resulted from the pandemic will require input from stakeholders with complementary ethical, scientific, and socioeconomic perspectives. It is our hope that as a community, we can rise to the challenge.

## Supporting information

supplemental alignment

## Acknowledgements

We thank the people in the Primate Genome Variation Consortium and BGI Primate Reference Consortium for allowing us to publish partial gene information early due to the urgency for COVID-19 research. We thank all persons and groups who shared samples for this research, including many samples from the Duke Lemur Center. ADM is supported by the Natural Sciences and Engineering Research Council of Canada (NSERC Discovery Grant) and Canada Research Chairs Program. MCJ’s postdoctoral appointment is supported by funding from the Natural Environment Research Council (NERC NE/T000341/1). IG and MG acknowledge the support of the Spanish Ministry of Science and Innovation through the Instituto de Salud Carlos III and the 2014–2020 Smart Growth Operating Program, to the EMBL partnership and cofinancing with the European Regional Development Fund (MINECO/FEDER, BIO2015-71792-P). We also acknowledge the support of the Centro de Excelencia Severo Ochoa, and the Generalitat de Catalunya through the Departament de Salut, Departament d’Empresa i Coneixement and the CERCA Programme. TMB is supported by funding from the European Research Council (ERC) under the European Union’s Horizon 2020 research and innovation programme (grant agreement No. 864203), BFU2017-86471-P (MINECO/FEDER, UE), “Unidad de Excelencia María de Maeztu”, funded by the AEI (CEX2018-000792-M), Howard Hughes International Early Career, Obra Social “La Caixa” and Secretaria d’Universitats i Recerca and CERCA Programme del Departament d’Economia i Coneixement de la Generalitat de Catalunya (GRC 2017 SGR 880). PSA thanks the National Institutes of Health (R35GM130333) for financial support. E.L is supported by CGL2017-82654-P (MINECO/FEDER,UE). EDJ and OF’s contributions were supported by funds from Howard Hughes Medical Institute and the Rockefeller University. Chris Smith drew the images for Figure 1. The authors would like to thank the Veterinary and Zoology staff at Wildlife Reserves Singapore for their help in obtaining the tissue samples, as well as the Lee Kong Chian Natural History Museum for storage and provision of the tissue samples.

## Author Contributions

ADM and JPH designed the study. ADM and JPH wrote the paper with input and edits from JDO, MCJ, LK, HR, JR, TMB, SEJ, and PSA. HR wrote the discussion on implications for lemur conservation in Madagascar. JDO and MCJ conducted genetic analyses with input from ADM. AV, LK, DJ. AN, LK and AN conducted gene sequence generation and alignment. Samples were provided by JEH, CR, ACK, CCK, WKL, JGHL, PT, and GU. JR, IG, MG, EL, RM, RAH and KK-HF. WD and GZ contributed to laboratory work, sequencing, genome reference assemblies and bioinformatics. FM ran the protein-protein interaction and substitution models with input from PSA. All authors have approved the final submission for publication.

## Data Availability Statement

All nucleotide and amino acid sequences used in this study are provided as supplemental text files.

## Figure Captions

**Supplementary Table S1:** Samples used to generate new ACE-2 sequences for this study

| Sample No | Infraorder | Family | Genus | Species |
| --- | --- | --- | --- | --- |
| 1 | Lemuroidea | Cheirogaleidae | *Cheirogaleus* | *major* |
| 2 | Lemuroidea | Cheirogaleidae | *Cheirogaleus* | *medius* |
| 3 | Lemuroidea | Cheirogaleidae | *Microcebus* | *murinus* |
| 4 | Lemuroidea | Cheirogaleidae | *Mirza* | *zaza* |
| 5 | Lemuroidea | Cheirogaleidae | *Mirza* | *zaza* |
| 6 | Lemuroidea | Daubentoniidae | *Daubentonia* | *madagascariensis* |
| 7 | Lemuroidea | Indriidae | *Avahi* | *laniger* |
| 8 | Lemuroidea | Indriidae | *Avahi* | *peyrierasi* |
| 9 | Lemuroidea | Indriidae | *Propithecus* | *coquereli* |
| 10 | Lemuroidea | Indriidae | *Propithecus* | *coronatus* |
| 11 | Lemuroidea | Indriidae | *Propithecus* | *diadema* |
| 12 | Lemuroidea | Indriidae | *Propithecus* | *diadema* |
| 13 | Lemuroidea | Indriidae | *Propithecus* | *edwardsi* |
| 14 | Lemuroidea | Indriidae | *Propithecus* | *perrieri* |
| 15 | Lemuroidea | Indriidae | *Propithecus* | *perrieri* |
| 16 | Lemuroidea | Indriidae | *Propithecus* | *tattersalli* |
| 17 | Lemuroidea | Indriidae | *Propithecus* | *verreauxi* |
| 18 | Lemuroidea | Lemuridae | *Eulemur* | *albifrons* |
| 19 | Lemuroidea | Lemuridae | *Eulemur* | *collaris* |
| 20 | Lemuroidea | Lemuridae | *Eulemur* | *collaris* |
| 21 | Lemuroidea | Lemuridae | *Eulemur* | *collaris* |
| 22 | Lemuroidea | Lemuridae | *Eulemur* | *coronatus* |
| 23 | Lemuroidea | Lemuridae | *Eulemur* | *coronatus* |
| 24 | Lemuroidea | Lemuridae | *Eulemur* | *flavifrons* |
| 25 | Lemuroidea | Lemuridae | *Eulemur* | *flavifrons* |
| 26 | Lemuroidea | Lemuridae | *Eulemur* | *fulvus* |
| 27 | Lemuroidea | Lemuridae | *Eulemur* | *macaco* |
| 28 | Lemuroidea | Lemuridae | *Eulemur* | *macaco* |
| 29 | Lemuroidea | Lemuridae | *Eulemur* | *mongoz* |
| 30 | Lemuroidea | Lemuridae | *Eulemur* | *rubriventer* |
| 31 | Lemuroidea | Lemuridae | *Eulemur* | *rufus* |
| 32 | Lemuroidea | Lemuridae | *Eulemur* | *sanfordi* |
| 33 | Lemuroidea | Lemuridae | *Eulemur* | *sanfordi* |
| 34 | Lemuroidea | Lemuridae | *Hapalemur* | *alaotrensis* |
| 35 | Lemuroidea | Lemuridae | *Hapalemur* | *gilberti* |
| 36 | Lemuroidea | Lemuridae | *Hapalemur* | *griseus* |
| 37 | Lemuroidea | Lemuridae | *Hapalemur* | *meridionalis* |
| 38 | Lemuroidea | Lemuridae | *Hapalemur* | *occidentalis* |
| 39 | Lemuroidea | Lemuridae | *Lemur* | *catta* |
| 40 | Lemuroidea | Lemuridae | *Prolemur* | *simus* |
| 41 | Lemuroidea | Lemuridae | *Varceia* | *variegata* |
| 42 | Lemuroidea | Lemuridae | *Varecia* | *rubra* |
| 43 | Lemuroidea | Lepilemuridae | *Lepilemur* | *ankaranensis* |
| 44 | Lemuroidea | Lepilemuridae | *Lepilemur* | *ankaranensis* |
| 45 | Lemuroidea | Lepilemuridae | *Lepilemur* | *dorsalis* |
| 46 | Lemuroidea | Lepilemuridae | *Lepilemur* | *ruficaudatus* |
| 47 | Lemuroidea | Lepilemuridae | *Lepilemur* | *septentrionalis* |
| 48 | Lemuroidea | Lepilemuridae | *Lepilemur* | *septentrionalis* |
| 49 | Lorisoidea | Galagidae | *Galago* | *moholi* |
| 50 | Lorisoidea | Galagidae | *Galago* | *senegalensis* |
| 51 | Lorisoidea | Galagidae | *Galagoides* | *demidovii* |
| 52 | Lorisoidea | Galagidae | *Otolemur* | *crassicaudatus* |
| 53 | Lorisoidea | Galagidae | *Otolemur* | *garnettii* |
| 54 | Lorisoidea | Galagidae | *Otolemur* | *monteiri* |
| 55 | Lorisoidea | Lorisidae | *Arctocebus* | *calabarensis* |
| 56 | Lorisoidea | Lorisidae | *Loris* | *lydekkerianus* |
| 57 | Lorisoidea | Lorisidae | *Loris* | *lydekkerianus* |
| 58 | Lorisoidea | Lorisidae | *Loris* | *tardigradus* |
| 59 | Lorisoidea | Lorisidae | *Nycticebus* | *bengalensis* |
| 60 | Lorisoidea | Lorisidae | *Nycticebus* | *bengalensis* |
| 61 | Lorisoidea | Lorisidae | *Nycticebus* | *bengalensis* |
| 62 | Lorisoidea | Lorisidae | *Nycticebus* | *bengalensis* |
| 63 | Lorisoidea | Lorisidae | *Nycticebus* | *pygmaeus* |
| 64 | Lorisoidea | Lorisidae | *Nycticebus* | *pygmaeus* |
| 65 | Lorisoidea | Lorisidae | *Perodicticus* | *ibeanus* |
| 66 | Lorisoidea | Lorisidae | *Perodicticus* | *potto* |

**Supplementary Table S2.** Results of alanine scanning mutagenesis experiments predicting critical binding sites between ACE2 and Sars-CoV-2 receptor binding domain. Residues whose mutation to alanine decrease the binding energy by ΔΔG_bind_ ≥1.0 kcal/mol are considered to be significant for binding (in bold).

| Human Residue | $\Delta\Delta G$ (kcal/mol)<sup>†</sup> |
| --- | --- |
| **Y41** | **4.1** |
| **D355** | **3.5** |
| **Q42** | **2.3** |
| **Y83** | **2.1** |
| **R357** | **2** |
| **D38** | **1.4** |
| **E37** | **1.2** |
| **Q24** | **1.1** |
| **K353** | **1.1** |
| T27 | 0.7 |
| H34 | 0.7 |
| K31 | 0.6 |
| D30 | 0.6 |
| E35 | 0.5 |
| L79 | 0.5 |
| L45 | 0.5 |
| F28 | 0.3 |
| M82 | 0.2 |
| N330 | 0.2 |
| L351 | 0.03 |

<sup>†</sup>The computational alanine mutagenesis analysis was performed with Rosetta and PDB file 6M0J.

